# Structural basis of ligand selectivity by a bacterial adhesin lectin involved in multi- species biofilm formation

**DOI:** 10.1101/2020.11.18.389155

**Authors:** Shuaiqi Guo, Tyler D.R. Vance, Hossein Zahiri, Robert Eves, Corey Stevens, Jan-Hendrik Hehemann, Silvia Vidal-Melgosa, Peter L. Davies

**Author notes:** Department of Laboratory Medicine and Pathobiology, University of Toronto, Toronto, ON Canada, M5S 1A8. Laboratoire des Polymères, Institut des Matériaux and Institut des Sciences et Ingénierie Chimiques, École Polytechnique Fédérale de Lausanne (EPFL), Batiment MXD, Station 12, 1015 Lausanne, Switzerland. Corresponding author: Peter L. Davies, **Email:**.

## Abstract

Carbohydrate recognition by lectins governs critical host-microbe interactions. *MpPA14* lectin is a domain of a 1.5-MDa adhesin responsible for a symbiotic bacterium-diatom interaction in Antarctica. Here we show *Mp*PA14 binds various monosaccharides, with L-fucose and N-acetyl glucosamine being the strongest ligands (K_d_ ~ 150 μM). High-resolution structures of *Mp*PA14 with 15 different sugars bound elucidated the molecular basis for the lectin’s apparent binding promiscuity but underlying selectivity. *Mp*PA14 mediates strong Ca^2+^-dependent interactions with the 3, 4 diols of L-fucopyranose and glucopyranoses, and binds other sugars via their specific minor isomers. Thus, *Mp*PA14 only binds polysaccharides like branched glucans and fucoidans with these free end-groups. Consistent with our findings, adhesion of *Mp*PA14 to diatom cells was selectively blocked by L-fucose, but not by N-acetyl galactosamine. With *Mp*PA14 lectin homologs present in adhesins of several pathogens, our work gives insight into an anti-adhesion strategy to block infection via ligand-based antagonists.

## Introduction

Carbohydrate-based polymers (glycans) are abundant on the exterior of cells(1,2). The recognition of glycans by carbohydrate-binding proteins or lectins, underlies many essential biological events such as fertilization, immunological responses, cell-to-cell communications, and host-pathogen interactions(3–9). Some lectins are components of larger adhesion proteins (adhesins), which are key virulence factors that mediate the attachment of bacteria to host cells and other surfaces(6, 10–12). Adhesins further mediate the colonization of bacteria in their favorable niches by helping develop biofilms, which cause over 80% of chronic infections in humans. Once formed, biofilms are resistant to various bactericidal treatments, including the use of antibiotics. With a current shortage of methods to treat biofilm-related diseases and the emerging prevalence of antibiotic-resistant bacteria, there is an urgent need to develop new strategies to treat bacterial infections. Since adhesins play a critical role in the initial stages of biofilm formation, the development of adhesin lectin antagonists holds great promise for treating various infections by blocking bacterial adhesion to human cells. To date, this anti-adhesion strategy has led to the development of some promising treatments against diseases. For example, binding of uropathogenic *Escherichia coli* (UPEC) to mannose-containing glycoproteins of human uroepithelium via the adhesin lectin FimH is an enabling step towards most urinary tract infections(13, 14). Mannoside-based FimH antagonists developed through structure-guided design can effectively block UPECs from binding to the human uroepithelium(15, 16). These compounds have demonstrated fast-acting efficacy against chronic urinary tract infections, and can prevent the disease when used as prophylactics(17–19). Despite these successes, widespread application of this anti-adhesion approach to treat other bacterial infections is hampered by a lack of knowledge at the molecular level of ligand recognition by other adhesin lectin modules.

*Marinomonas primoryensis* ice-binding protein (*Mp*IBP) is a large (~ 1.5 MDa) Repeats-In-Toxin (RTX) adhesin found on the surface of its Antarctic Gram-negative bacterium(11,20–22). While the N terminus of *Mp*IBP anchors the giant protein to the bacterial outer membrane, the ligandbinding modules near the C terminus bind the bacterium to both ice and photosynthetic diatoms to form symbiotic biofilms on the underside of sea ice. *Mp*IBP was initially extracted from the cell lysate of its native bacterium by an ice-affinity purification step(20). Intriguingly, the protein failed to elute from a Superdex S200 size-exclusion column intended for further purification. This suggested that the adhesin interacts with the Superdex matrix, which is based on porous agarose particles covalently linked to dextran, a complex branched polymer of α-D-1,6-glucose. Bioinformatic analyses indicated the presence of a ~ 20-kDa domain near the C terminus of *Mp*IBP that is a member of the PA14 family, which is a carbohydrate-binding lectin module widely distributed across several kingdoms of life(23, 24). PA14 homologues are found in human proteins like fibrocystin(25), and fungal and bacterial proteins such as β-glucosidases(26, 27), proteases(23), and adhesins(10, 21,28, 29). PA14 domains share a common β-sandwich fold and the presence of two consecutive aspartate residues in a *cis* peptide linkage (D*cis*D). The D*cis*D motif coordinates a Ca^2+^ ion that is directly involved in binding polar vicinal hydroxyl groups of various carbohydrates(10, 12, 29). Despite these conserved features in their ligand-binding sites, PA14 lectins in microbial adhesins have a broad specificity profile for a range of carbohydrates. In this regard, how PA14 lectins of bacterial adhesins recognize their ligands remains unclear. Yet, this highly conserved module is widespread in adhesins of many different bacteria, including those of human pathogens. These important attributes justify the pursuit of structural studies to elucidate the molecular basis of ligand recognition by *MpPA14,* which may set the stage for the subsequent development of antagonists to block harmful adhesion.

In this report, we use various types of binding assays and glycan microarrays to show that *MpPA14* is a lectin with an unusual binding promiscuity to monosaccharides but is specific in binding certain polysaccharides. X-ray crystal structures of *Mp*PA14 in complex with 15 different simple sugars at atomic resolution reveal the molecular basis for the uncommon ligand selectivity by the lectin. We further show that the adhesion of *Mp*PA14 to its host diatom cells can be fully abolished by a micromolar concentration of L-fucose. Since bioinformatic analyses indicate that lectins highly similar to *Mp*PA14 are present in many bacterial adhesins, including those from human pathogens such as *Vibrio cholerae* and *Vibrio vulnificus* (12), there is an opportunity to use a structure-based approach to devise high-affinity lectin antagonists to block harmful biofilm formation or to develop molecular probes to detect these microbes.

## Results and Discussion

### MpPA14 interacts strongly with fucose and N-acetyl glucosamine

To gain insight into the binding specificity of *Mp*PA14, we investigated the relative affinity of various monosaccharides for the lectin by a comparative competition assay(12). *Mp*PA14 bound to Superdex resin was competitively released into solution by the progressive addition of free sugars. The released protein concentrations measured by absorbance at 280 nm were plotted as a function of free sugar concentration to produce semi-quantitative binding curves (Figure 1A, B). The apparent dissociation constant (K_d_app) calculated from these binding curves was used as a relative measure of affinity for each assayed sugar (Table 1). *Mp*PA14 lectin bound L-fucose the strongest with a K_d_app of 0.65 mM, followed by GlcNAc (K_d_app 1.07 mM; Figure 1A). Glucose and 2-deoxy-glucose bound the lectin with similar affinity, giving K_d_app values of 1.36 and 1.37 mM, respectively. D-mannose, and methyl-α-D-glucose bound *Mp*PA14 with slightly weaker affinity (K_d_app = 1.7 mM and 2.1 mM, respectively), and the binding of D-allose, 3-O-methyl-D-glucose and D-galactose was significantly diminished to K_d_app values between 4.1 and 6.8 mM. There was no measurable interaction between *Mp*PA14 and N-acetyl-galactosamine (GalNAc). D-ribose bound dextran more strongly than its derivative 2-deoxy-D-ribose (K_d_app of 6.8 mM as opposed to 18 mM; Figure 1B, Table 1). L-arabinose exhibited higher affinity for *Mp*PA14 than the other pentoses, with a K_d_app of 2.2 mM.

**Figure 1:**
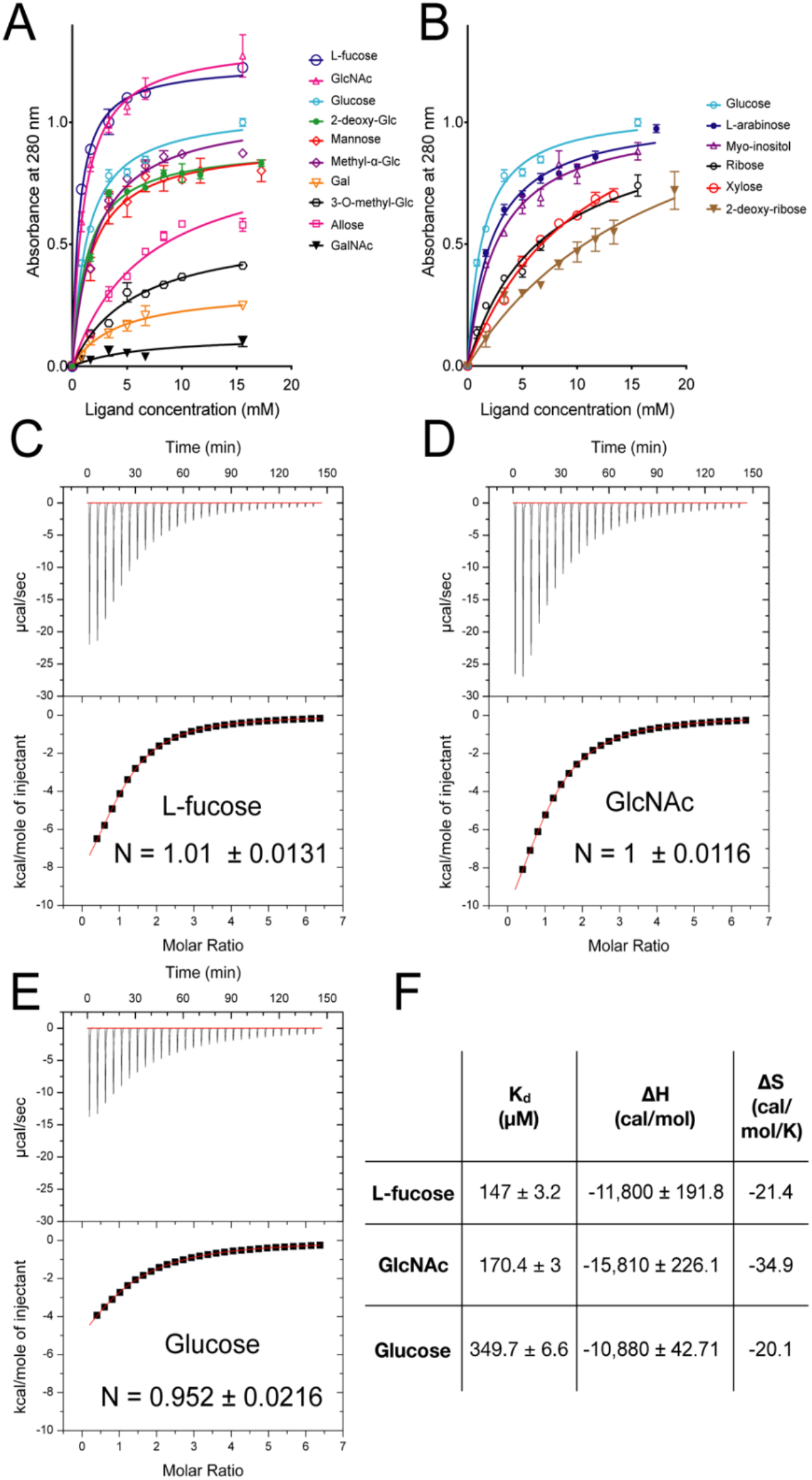
Competition binding assay to determine relative affinity of *Mp*PA14 for simple carbohydrates. A) Dextran-based competition binding assay to determine relative affinity of *Mp*PA14 for various hexoses and their derivatives. Each experiment was done in triplicate. B) Dextran-based competition binding assay to determine relative affinity of *Mp*PA14 for selected pentoses and myo-inositol. Isothermal titration calorimetry of *Mp*PA14 binding to fucose (C), GlcNAc (D), and glucose (E), respectively. The binding stoichiometry (N) is indicated on each graph. (F) Table of the calculated thermodynamic parameters, including the dissociation constant (Kd), enthalpy (ΔH) and entropy (ΔS) for each of the three sugars.

**Table 1.** Apparent dissociation constants for the binding of different sugars.

| Sugars | K<sub>D</sub>app (mM) |
| --- | --- |
| L-fucose | 0.65 |
| N-acetyl-glucosamine | 1.07 |
| Glucose | 1.4 |
| 2-deoxy-glucose | 1.4 |
| Mannose | 1.7 |
| Methyl-α-glucose | 2.1 |
| L-arabinose | 2.2 |
| Inositol | 2.7 |
| Galactose | 4.1 |
| 3-*O*-methyl-glucose | 5.7 |
| Allose | 6.8 |
| Ribose | 6.8 |
| Xylose | 11 |
| 2-deoxy-ribose | 18 |
| N-acetyl-galactosamine | - |

To further assess the binding thermodynamic parameters of *Mp*PA14 ligands, we performed isothermal titration calorimetry (ITC) with L-fucose, N-acetyl-glucosamine (GlcNAc) and glucose (Figure 1C-E). The ITC measurements produced rectangular hyperbolic curves for all three simple saccharides, and fitted well to a one binding-site model with calculated N values being close to 1 ^(12, 21)^. ITC ranked the affinity of these three ligands in the same order shown by the competition binding assay, with L-fucose as the strongest ligand followed by GlcNAc and then glucose. The K_d_ values calculated from the ITC measurements were significantly lower than the K_d_app values obtained from the competition assay (Figure 1F, Table 1). However, this is to be expected because the dextran beads used in the competition assay have multiple binding sites nearby that attract the lectin, whereas the calorimetry was done with free sugars in solution. With K_d_ values of 147 μM and 170 μM for L-fucose and GlcNAc, these two ligands had a greater than 2-fold higher affinity for *Mp*PA14 than did glucose (K_d_ = 350 μM). In general, the affinity of lectins for monosaccharides (K_d_ values) lies within the high micromolar to millimolar range(8, 30, 31). Thus, our results showed the *Mp*PA14 domain from its bacterial adhesin had relatively high affinity for the three strongest ligands in comparison to other lectins. Furthermore, negative enthalpic (ΔH) and entropic (ΔS) contributions were calculated for all three carbohydrates when they bound to *Mp*PA14, which indicated the binding was driven primarily by polar interaction such as the formation of hydrogen and ionic bonds rather than hydrophobic interactions. This was consistent with the observation that the Ca^2+^-dependent ligand-binding site of *Mp*PA14 consisted of mainly polar and charged amino acids without any residues with large hydrophobic side chains. We therefore acquired detailed structural information to study *MpPA14* ligand recognition.

### Structural basis of MpPA14 selectivity for glucopyranoses

To examine the molecular basis of carbohydrate recognition by *Mp*PA14, we determined the X-ray crystal structures of the lectin in complex with 14 new saccharides to a resolution of 1-1.3 Å (Table S1-3). The lectin fold is a β-sandwich domain that binds four to seven Ca^2+^ ions (Ca1-7) on its surface(21) (Figure S1A). Ca1 is coordinated by the D*cis*D motif (Asp110 and Asp111) on the periphery of the protein, which is directly involved in binding carbohydrate with help from amino-acid residues in loops 9 and 11 (L9 and L11; Figure S1B). While Ca 2-4 likely have a role in lectin folding(12, 21) (Figure S1C-E), Ca5-7 have few ligands from the lectin and are probably an artifact from the crystallization condition that contained over 100 mM of CaCl_2_. There were no substantial conformational changes to the overall lectin fold when it was complexed with different sugars (r. s. m. d. <0.1 Å).

As suggested by ITC, sugar recognition by *Mp*PA14 is primarily driven by polar interactions in a Ca^2+^-dependent manner. Glucose, GlcNAc, and other glucopyranose-containing carbohydrates including 2-deoxy-glucose, methyl-α-glucose and two disaccharides, sucrose and trehalose, all bound to the *Mp*-PA14-Ca1 via their 3, 4 *trans* vicinal diols in gauche configuration with a dihedral angle of ~ 60° (Figure 2A, Figure S2, S3A, B). This interaction is further enhanced with the diol being coordinated by the side-chain carboxyl and hydroxyl oxygens of the D*cis*D motif, and main-chain and side-chain protein ligands from L11 (Gln156, Gly157 and Asp159, Figure 2B). The acetyl group on the C-2 position of GlcNAc interacts with the side-chain atoms of Asp 159 on L11 (Figure 2C), holding the aspartate side chain down in one stable conformation. This additional interaction likely accounts for the higher affinity of GlcNAc to the lectin than for other glucopyranoses.

**Figure 2:**
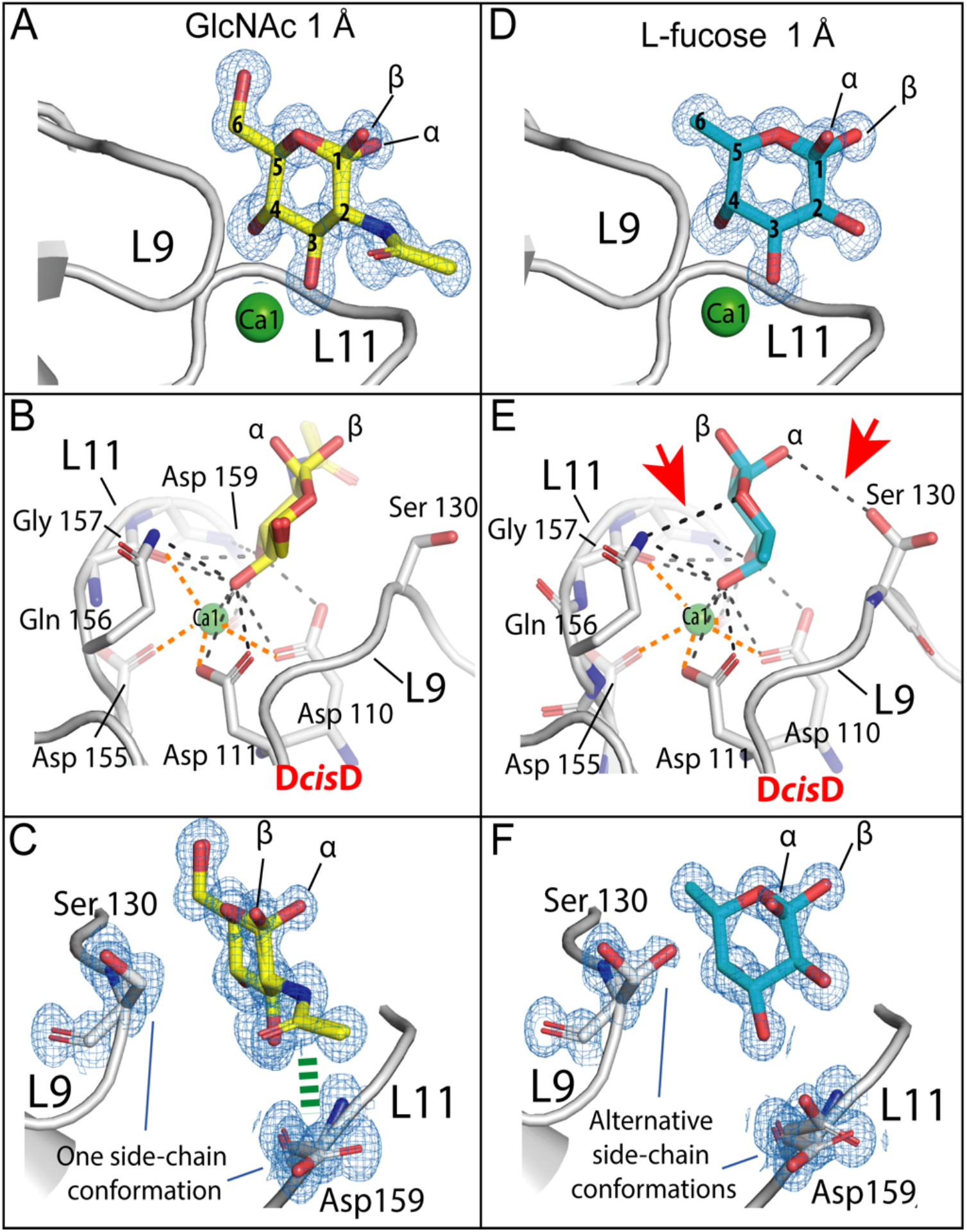
Ligand-binding site of *Mp*PA14 in complex with GlcNAc and L-fucose. Structural features of the *Mp*PA14 ligand-binding sites with GlcNAc (A) and L-fucose (D). Detailed carbohydrate-protein interactions between *Mp*PA14 and GlcNAc (B), and L-fucose (E), respectively. Ionic bonds between Ca1 and *Mp*PA14 are indicated by thicker orange dashed lines, while polar interactions between the lectin and carbohydrates are indicated with black dashed lines. Red arrowheads in (E) indicate the additional polar interactions between L-fucose and *Mp*PA14 compared to those in (B). (C, F) Angled view comparing the conformations of the side chains of L9-Ser130, and L11-Asp159 in the ligand-binding sites of *Mp*PA14-GlcNAc (C) and *Mp*PA14-fucose (F) complexes. Carbon atoms of GlcNAc are colored yellow, while those of L-fucose and *Mp*PA14 are colored cyan and light grey, respectively. The interaction between the acetyl group of GlcNAc and side chain of Asp159 is indicated by a green dashed line. Oxygens are colored red, nitrogens are colored blue and Ca^2+^ ions are colored green. The 2 F_o_ – F_c_ maps shown for sugars and ligand-binding site residues in (A), (C), (D) and (F) are shown as blue meshes (contoured at σ=1).

Structural data shown here for *Mp*PA14 contrast with the previously reported structure of *Mh*PA14 in complex with glucose(12). *Mh*PA14 is a *Mp*PA14 homolog from an RTX adhesin of the oil-degrading bacterium, *Marinobacter hydrocarbonoclasticus.* With a similar ligand-binding site to *Mp*PA14(12), *Mh*PA14 also had a strong preference for binding L-fucose and glucopyranoses over other monosaccharides. However, X-ray crystallography showed *Mh*PA14 complexing glucopyranose via its 1,2 diol. Given that the C-2 position of GlcNAc lacks the hydroxyl group required for interacting with *Mp*PA14 via the 1,2 diol, the binding mode shown by the *Mh*PA14-glucose complex could not explain the lectin’s high affinity for this acetylated sugar. Close inspection of the *Mh*PA14-glucose complex structure revealed that the monosaccharide in the carbohydrate-binding site is in direct contact with a neighboring symmetry-related molecule, indicating the tight packing of the unit cell might have caused the sugar to bind in a less favorable configuration. Moreover, the *apo-Mh*PA14 structure showed that its unoccupied ligand-binding site is involved in crystal contact with the symmetry-related molecules, leaving insufficient space for carbohydrate binding. The observed crystal-packing artifacts explained why co-crystallization of *Mh*PA14 with various other sugars, and direct soaking experiments with the apo-*Mh*PA14 crystal were futile.

### Why L-fucose is a better ligand than glucopyranoses

Based on results from docking experiments, it was proposed that L-fucose binds *Mh*PA14 via its 2,3 diol(12). However, the well-resolved 1-Å electron density map in this study unambiguously showed fucose bound *Mp*PA14 with its *cis* 3, 4 diol in the gauche conformation with a dihedral angle of 47° (Figure 2D). In contrast to the *Mp*PA14 hexose ligands, which are all in the D-configuration, fucose is in the L-configuration, with hydroxyl groups of the fucopyranose ring pointed in opposite directions. This helps the endocyclic oxygen atom of fucose to point toward L11 and hydrogen bond with the sidechain of Gln156 (Figure 2E). Additionally, the hydroxyl on the α-anomeric carbon may hydrogen bond with the sidechain of Ser130 on L9, clamping the pyranose ring tightly into the binding site (Figure 2E, F). The L-fucose-*Mp*PA14 interaction is distinct from that shown by the bacterial C-type lectin, LecB, from *Pseudomonas aeruginosa,* which uses two side-by-side Ca^2+^ ions to directly coordinate the 2, 3, 4 triol of L-fucose(32, 33). The additional ionic interaction between LecB and L-fucose can explain its enhanced affinity (K_d_ = 58 μM) for the sugar compared to that of *Mp*PA14 (K_d_ = 147 μM).

### Promiscuity of MpPA14 in monosaccharide recognition

To investigate the molecular basis for the plastic nature of *Mp*PA14 in binding various monosaccharides, we investigated *Mp*PA14 structures in complex with various glucose epimers and derivatives.

Mannose bound *Mp*PA14 slightly weaker than glucose (Figure 1A, Table 1). The electron density map for the mannose-*Mp*PA14 complex indicated the sugar bound in two distinct conformations (Figure 3A, B, Figure S3B). Like glucose, D-mannose bound *Mp*PA14 via the 3, 4 diol. However, since the mannose C-2 hydroxyl moiety is in an axial position, its oxygen atom may clash with the β-carbon of the Ser130 as they are only 3 Å away from each other (Figure 3C). In addition, as the C-2 and C-3 hydroxyl groups of mannose are positioned in *cis*, they may form an intramolecular hydrogen bond that further weakens the 3, 4-diol from binding Ca1. Alternatively, β-mannopyranose can bind *Mp*PA14 using a second configuration where its 2,3 diol anchors the saccharide ring in an inverted fashion, allowing the ring oxygen to hydrogen bond with the side-chain amide group of the Gln156 (Figure 3D). However, α-mannopyranose failed to fit into the electron density via this binding mode, which indicates that *Mp*PA14 can only recognize the less prevalent β-anomer in the equilibrium via its 2,3 diol (33% β-mannopyranose as opposed to 62% α-mannopyranose at 30 °C)(34). This apparent lack of one distinct stable binding configuration explains the relatively inferior affinity of mannose compared to glucose.

**Figure 3:**
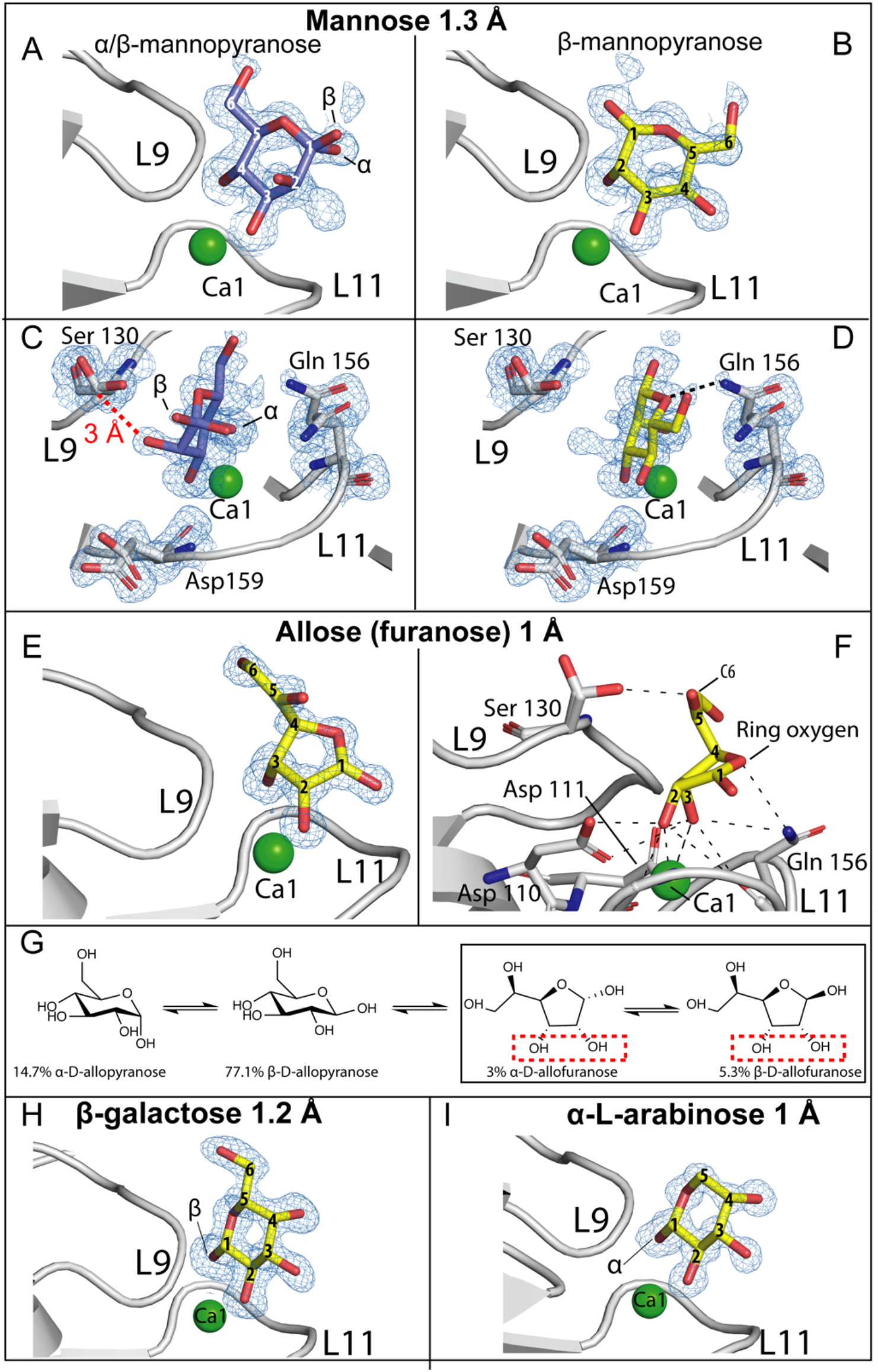
Ligand-binding site of *Mp*PA14 in complex with mannose, allose, galactose and L-arabinose. Side (A) and top views (C) of *a-* and β-mannopyranose rings anchored to *Mp*PA14 via their 3, 4 diols. Electron density for only β-mannopyranose was seen for the binding mode via the 2, 3 diol (side view in B and top-down view in D). The distance between the β-carbon of serine on L9 and C-2 hydroxyl oxygen of α/ β-mannopyranose is indicated by a red dashed line. E) Sideview of β-allofuranose in the *Mp*PA14 ligand-binding site. F) Detailed polar interactions (black dashed lines) between β-allofuranose and *Mp*PA14. Amino-acid residues involved in allose-*Mp*PA14 interaction are labelled. The color scheme is the same as in Fig. 3. (G) Equilibria of allose anomers in aqueous solution at 30 °C. The diol of allofuranose responsible for binding *Mp*PA14 is indicated by a red-dashed box. Side-view of β-galactose (H) and α-L-arabinose (I) in the *Mp*PA14 ligand-binding site.

With the C-3 hydroxyl in the axial position, allopyranose cannot bind *Mp*PA14 via its *cis* vicinal 3, 4 diol as this would cause the saccharide ring to clash with residues of L11 (e.g. Gln156). Unexpectedly, the binding site of the lectin contained the furanose form of allose with its 2,3 diol interacting with Ca1 (Figure 3E, Figure S3C). The furanose rings lean towards L11, with their endocyclic oxygen interacting with the side-chain amide group of Gln156, and the C-5 hydroxyl group extends out to hydrogen bond with the side-chain hydroxyl of Ser130 (Figure 3F). At 30°C, approximately 92% of allose exists as pyranoses in solution (Figure 3G). Yet *Mp*PA14 binding appears to be dependent on the rare presence of the allofuranose, which only makes up ~8% of allose in solution, explaining the feeble affinity of this sugar for the lectin (Table 1). Similarly, 3-0-methyl-glucose cannot bind *Mp*PA14 via the 3, 4 diol because of the substituted methyl group on its C-3. This sugar compensates by binding using the 1,2 diol of the β-anomer (Figure S4A), resulting in its considerably weaker binding to the lectin compared to glucopyranoses and mannose.

Galactose has its C-4 hydroxyl in the axial position instead of being equatorial as in glucose. Docking of the galactopyranose ring to *Mp*PA14 via the 3, 4 diol is not possible due to steric hinderance against Gln129 and Ser130 on L-9. Instead, galactose can only interact with *Mp*PA14 via the 1,2 diol of its rare β-anomer (Figure 3H). These limitations explain the relatively weak interaction between galactose and *Mp*PA14 (Table 1). Results from the binding analysis of GalNAc verify this assessment. Having just shown the 3, 4 diol of galactopyranose cannot complex *Mp*PA14, binding of GalNAc is completely abolished because its 1,2 diol is unavailable due to the C-2 hydroxyl being substituted with an acetyl group.

We further analyzed the crystal structures of *Mp*PA14 in complex with three pentoses: L-arabinose, ribose and 2-deoxy-ribose, as well as inositol, which has an unusual 6-carbon saccharide ring without an endocyclic oxygen (Figure S3B, C, S4). These four carbohydrates bound *Mp*PA14 weaker than glucose and mannose (Table 1). Crystal structures showed that *Mp*PA14 selects the pyranose form of pentoses for binding. As pentoses have a higher percentage of furanose present in the conformational equilibria than do hexoses, selectivity for the more thermodynamically stable pyranoses might contribute to the weaker affinity of pentoses towards the lectin. For instance, despite L-arabinose existing in solution predominantly in the pyranose form at 25 °C (57 % *a*-vs 30.5% β-arabinopyranose) (35), only the a-anomer complexes *Mp*PA14 via its 1,2 diol (Figure 3I). The same rationale can be used to explain the even weaker affinity of ribose and its derivative 2-deoxy ribose for *Mp*PA14 (Figure S4B, C) (36). In the case of inositol, the composite electron density map indicates it binds *Mp*PA14 in several different conformations (Figure S4D-F). This promiscuous binding mode is indicative of a lack of one stable binding conformation, which might explain inositol’s moderate affinity to *Mp*PA14.

In summary, X-ray crystallography has elucidated the molecular basis of *Mp*PA14’s promiscuity in binding a range of monosaccharides. Remarkably, the lectin can discern favorable conformations of these monosaccharides from their non-binding anomers, even when the latter are much more prevalent in the equilibria (e.g. allofuranose as opposed allopyranose). Nevertheless, monovalent or simple carbohydrate oligomers are typically not the physiological targets of lectins(30). In the context of *Mp*PA14, it likely binds complex glycans or glyco-proteins present on the surfaces of microbes where the proximity effect of having numerous identical or similar end groups increases the avidity of the lectin interaction. In this way the sugar-binding activity of *Mp*PA14 can help to form biofilms. This prompted us to survey the lectin-sugar interactions from a broad spectrum of complex carbohydrates using glycan microarray technology.

### MpPA14 binds glucopyranose and fucose moieties of complex glycans

To dissect *Mp*PA14’s role in the formation of mixed-species biofilms, we probed two different microbial glycan microarrays for lectin binding partners(37). We first analyzed the binding of GFP-*Mp*PA14 to 16 different polysaccharides consisting primarily of glucans and mannans from bacteria and fungi (Imperial College Glycosciences Laboratory). Four glucans: pullulan, lentinan, dextran and grifolan bound most avidly to *Mp*PA14 (Figure 4A). To emphasize the specificity of this interaction, six of the other polysaccharides showed negligible or no detectable binding, which included glucans such as curdlan and those purified from oat and barley, as well as mannoprotein, glucurono-xylomannan, and GN6-AO which is hexasaccharide of 1,4 linked GlcNAc (Chitin) with an aminooxy group (AO; Table S4). Consistent with findings from our structural and binding analyses (Figure 1–3), the four strongest binders contain multiple glucopyranoses with unoccupied 3, 4 diols either as internal or terminal moieties in their linear backbone (pullulan) as well as in their branches (lentinan, dextran and grifolan; Figure 4A). In contrast, weak or negligible binding to *Mp*PA14 was demonstrated for linear glucans formed through 1,3 and 1,4 linkages such as β-glucans of oat and barley. Each of these linear polysaccharides has only one 3, 4 diol set from their terminal glucopyranose accessible for *Mp*PA14 binding, while all potential 3, 4 diols in the backbone are involved in the formation glycosidic bonds (Figure 4A). Similarly, GN6-AO interacted poorly with *Mp*PA14 because it too has only one free 3, 4 diol in the terminal GlcNAc. Furthermore, moderate lectin-glycan interactions were observed for the highly branched N-mannoprotein from the fungus *Candida albicans* (Figure 4A, orange bar, Table S4), while binding to glucurono-xylomannan and mannoprotein from the fungus *Aspergillus fumigatus* was negligible as these glycans lack the more favorable structural epitopes of 3, 4 diols on either glucopyranose or L-fucose.

**Figure 4:**
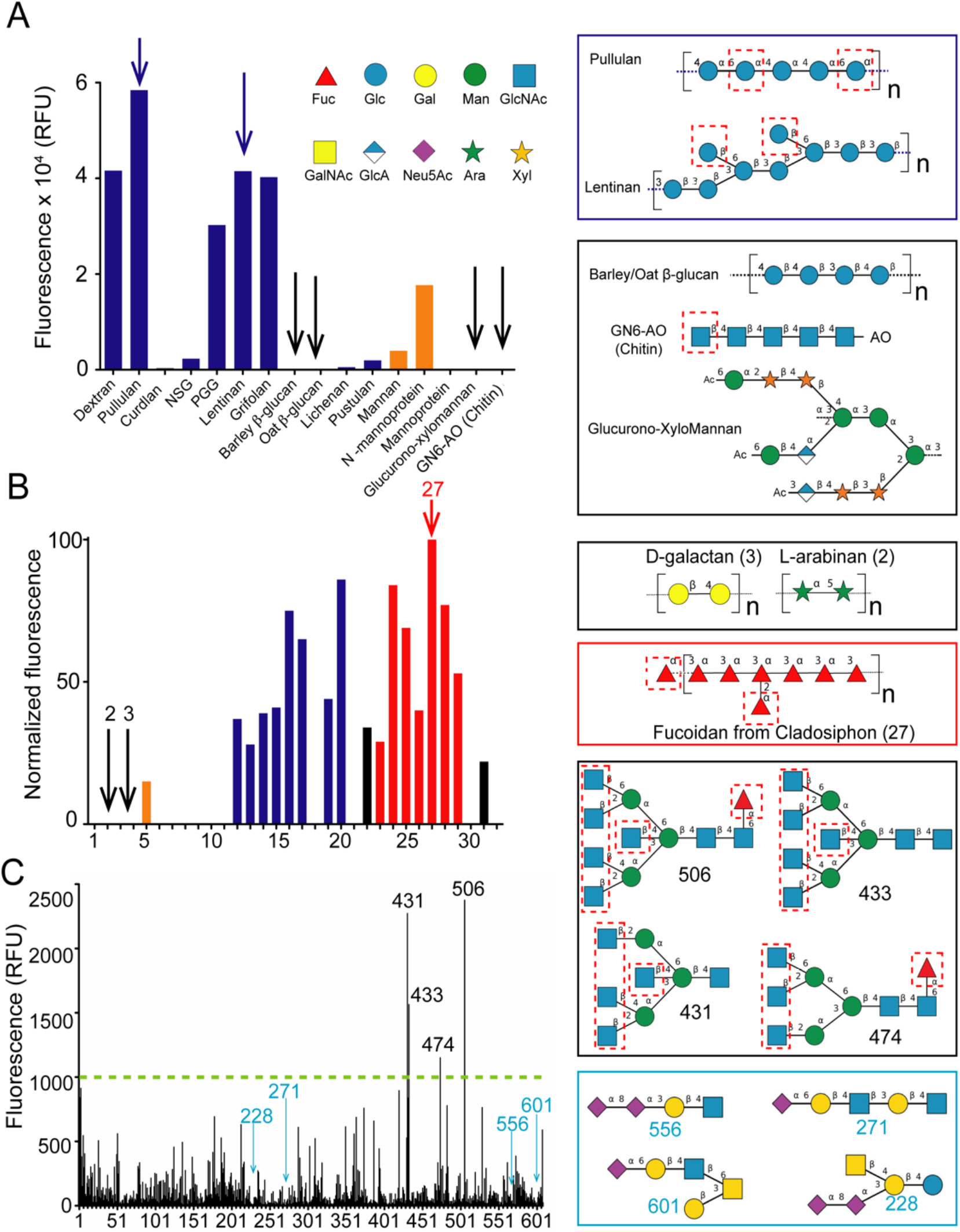
Binding of *Mp*PA14 to sugar oligomers identified by three different glycan microarrays. A) Microbial glycan array analysis conducted at Imperial College Glycosciences Laboratory. Data bars showing binding intensity for glucans and mannans are colored blue and orange, respectively. Examples of strong binders are drawn in the blue box (top right, blue arrows in the graph), while those for weak binders are drawn below (black arrows in the graph). The putative sites for *Mp*PA14 binding to the glycans are indicated by dashed red boxes. Glycan nomenclature symbols are included. Abbreviations are as follows: L-fucose, Fuc; Glucose, Glc; Galactose, Gal; Mannose, Man; N-acetyl-glucosamine, GlcNAc; N-acetyl-galactosamine, GalNAc; Glucuronic Acid, GlcA; N-acetyl-Neuraminic acid, Neu5Ac; L-arabinose, Ara; Xylose, Xyl. B) Microarray data of glycans from marine algae and land plants. Glucans (blue), mannans (orange) fucoidans (red), and other types of glycans (black). Examples of non-binders (right, black arrows in the graph), while the structure for Cladosiphon fucoidan is shown below (red arrow in the graph). C) Mammalian glycan array analyzed at the Consortium for Functional Glycomics. Representative structures for the strong *Mp*PA14 binders (black box) and non-binders (blue box) are drawn on the right.

**Figure 5:**
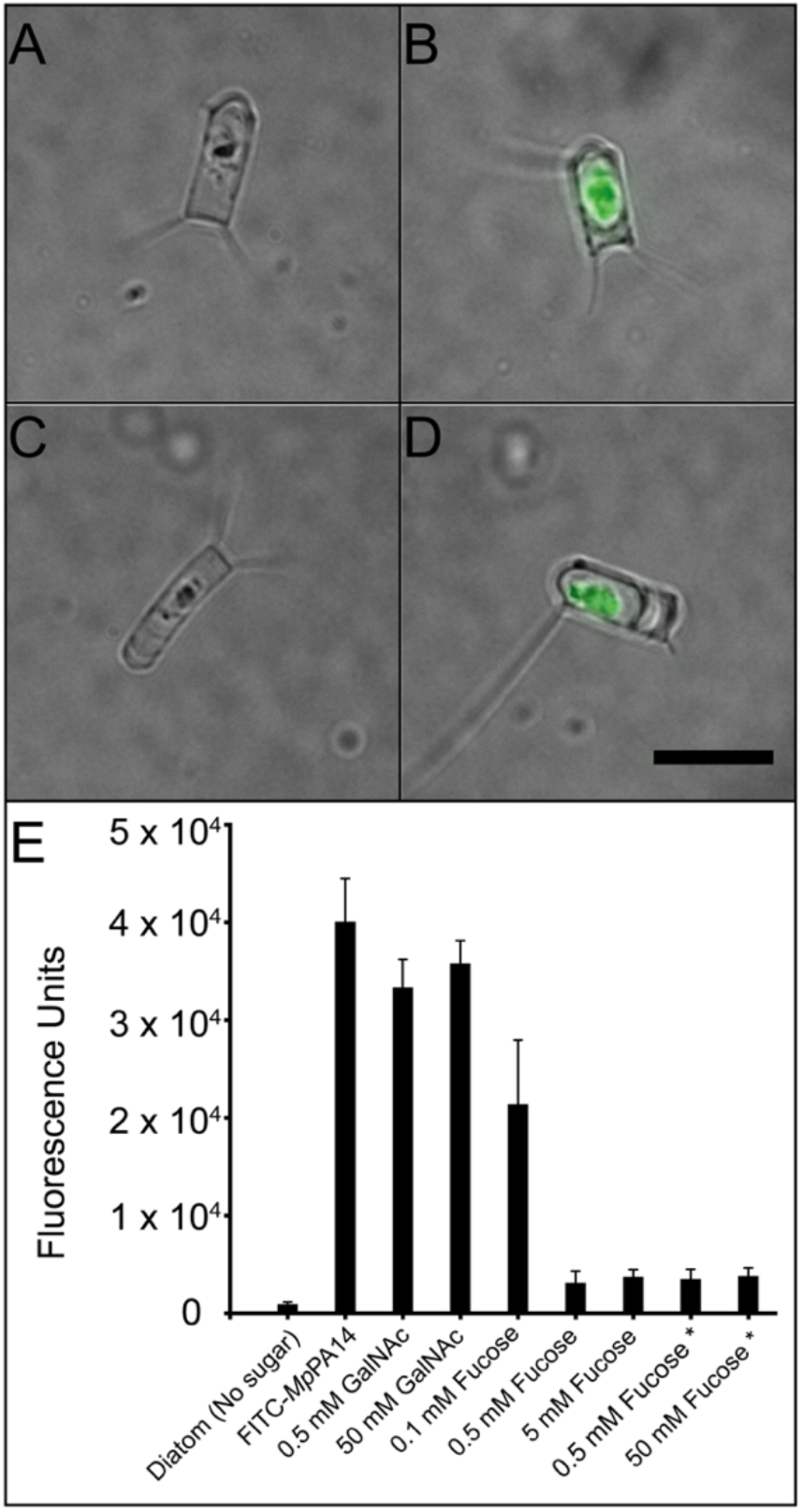
Inhibition of *Mp*PA14 binding to diatom *C. neogracile.* Representative images showing: (A) an untreated *C. neogracile* with basal autofluorescence in the center from its cell; (B) a *C. neogracile* cell treated with 0.2 mg/mL FITC-labelled *Mp*PA14: (C) a *C. neogracile* cell treated with 0.5 mM L-fucose and 0.2 mg/mL FITC-labelled *Mp*PA14; (D) a *C. neogracile* cell treated with 0.2 mg/mL FITC-labelled *Mp*PA14 and 50 mM GalNAc. All four images (A) – (D) are at the same scale, with the black scale bar in (D) indicating 10 μm (E) Fluorescence levels shown by the *C. neogracile* cells alone and those treated with various ligands. Each bar represents the quantification of average fluorescence from 30 individual diatoms. Data bars with an asterisk underneath represent experiments where diatoms were incubated with FITC-*Mp*PA14 before L-fucose was added.

To expand the repertoire of glycans beyond glucans and mannans, we performed a second focused microarray composed of 32 polysaccharides found in microbes such as fungi and bacteria, as well as macroalgae and plants (Figure 4B). Similar to the first microarray, *Mp*PA14 showed significant binding to some glucans that present multiple sets of unoccupied 3, 4 diols, which included pustulan, pachyman and scleroglucan (Figure 4B, dark blue bars; glycans 20, 16 and 17, respectively; Table S5). In addition, *Mp*PA14 interacted avidly with fucoidans from macroalgae. The strongest *Mp*PA14-binding fucoidans were to Cladosiphon(38), Sargassum and *Ascophyllum nodosum* (glycans 27, 24 and 28, red bars). Since fucoidans are algal polysaccharides primarily consisting of a linear backbone of sulphated a-1,3-or α-1,3;1,4-linked-L-fucose(39), *Mp*PA14 can interact by binding the terminal moieties on the backbone and branches with unoccupied 3, 4 diols. In contrast, *Mp*PA14 did not bind to polysaccharides such as arabinan, galactan, galactomannan, xylan, xyloglucan and porphyran (glycans 2, 3, 6, 11,21 and 32 respectively; Figure 4B) as these lack the structural epitopes required for favorable interactions with *Mp*PA14. For instance, while arabinan is a polymer of arabinose, a sugar that *Mp*PA14 binds, X-ray crystallography showed that *Mp*PA14 is selective in only forming a complex with a-L-arabinopyranose (Figure 3I). This conformer is not present in the arabinan polymer composed of 1,5 linked α-L-arabino-furanoses, which are then not free to transition to the pyranose form. Similarly, since the *Mp*PA14 binds galactose via the 1,2 diol (β-anomer) (Figure 3H), the lectin cannot interact with galactans as they are polymers of β-1,4-D-galactose (Figure 4B, Table S5).

Some differences were observed in the binding results with several polysaccharides present in both the first and second glycan microarrays. Examples include: mannan from *S. cerevisiae,* which showed weak binding to *Mp*PA14 in the first array (glycan number 12; Figure 4A), but had no detectable interaction with the lectin in the second array (glycan number 8; Figure 4B); Pullulan bound *Mp*PA14 more strongly than pustulan in the 1^st^ array (glycan numbers 2 and 11, respectively; Figure 4A), whereas their relative affinities were reversed in the 2^nd^ array (glycan numbers 12 and 20, respectively; Figure 4B). These minor discrepancies in the binding results could be due to differences in the polysaccharide sources or the different methods used to immobilize them onto the microarrays(40, 41). Nevertheless, both arrays pointed to the key result that *Mp*PA14 selectively binds glucans with multiple unoccupied 3, 4 diols, while the lectin does not recognize glycans such as arabinans, galactans and xylans. Furthermore, the finding that *Mp*PA14 interacted strongly with fucoidans is consistent with the *Mp*PA14-fucose interaction demonstrated by the binding and structural data. This result may have physiological relevance, as L-fucose-containing polysaccharides are highly prevalent in the exudates of diatoms(42), at least one of which is a natural host of *M. primoryensis.*

Since the PA14 domain is widespread in bacteria, including some that are human commensals and others that are pathogens, we reasoned that *Mp*PA14 and its homologs might interact with mammalian glycans. Therefore, we tested the lectin on a microarray consisting of 609 complex mammalian glycans (Consortium for Functional Glycomics, version 5.2). Four glycans (506, 431, 433 and 474) stood out amongst the strongest binders (Figure 4C). They share a common architecture as moderately branched mannose-containing oligomers with a bisecting GlcNAc motif. Each of the four glycans has three to five terminal GlcNAc moieties with 3, 4 diols available for complexing *Mp*PA14. Glycans 506 and 474 also have one α-L-fucose moiety linked to the surface-immobilized GlcNAc. Interestingly, the strongest binder, glycan 506 differs from glycan 433 only by the addition of the α-L-fucose, suggesting this monosaccharide might contribute to the higher affinity of glycan 506 by presenting an extra binding site for the lectin. Glycans with fewer binding epitopes of GlcNAc and fucose bound weaker in general. Out of the 61 glycans that did not bind *Mp*PA14 (RFUs of 20 or below), 33 have N-acetyl-neuraminic acid as their terminal sugar, while another 25 of these non-binders end with either galactose or GalNAc (Figure 4C). Indeed, as shown by our structural analyses above, these sugars lack the 3, 4 diol motifs of glucopyranoses and L-fucose preferred for *Mp*PA14 recognition.

### MpPA14 lectin homologs are found in pathogens

The binding conformations of various monosaccharides identified in the structural analyses can lay the foundation for the structure-guided design of glycan-based probes for detecting microbes or for making inhibitors to disrupt bacterial adhesion. *Mp*PA14 homologs are widespread in the adhesins of Gram-negative bacteria such as the previously reported *Mh*PA14 from the oil-degrading *M. hydrocarbonoclasticus,* as well as those adhesins that help pathogenic bacteria infect specific niches. For example, a large RTX adhesin from the flesh-eating human pathogen, *Vibrio vulnificus,* contains a *Mp*PA14 homolog (*Vv*PA14) with 41.5% identity at the protein level. Moreover, the amino-acid residues involved in coordinating Ca1 and recognizing glycans are conserved between the *Mp*PA14 and *Vv*PA14 (Figure S1F). Despite small deviations in aminoacid sequence between *Mp*PA14 and *Mh*PA14 from *M. hydrocarbonoclasticus*, these lectins have the same monosaccharide ligands and there is insignificant variation in their complex glycan recognition (12). With its ligand-binding site even more like *Mp*PA14 than that of *Mh*PA14, *Vv*PA14 probably binds to the same simple sugars. It is therefore of interest to test the inhibitory effect of strong PA14 binders identified in this study to set the stage for developing novel strategies for modulating bacterial adhesion.

### L-fucose blocks MpPA14-diatom interaction

Having identified L-fucose as the strongest monosaccharide ligand for *Mp*PA14, we set out to validate its potential as an inhibitor for the lectin-dependent bacteria-diatom interaction that led to the discovery and characterization of this protein(21). Here we tested if L-fucose can block fluorescently labelled *Mp*PA14 from binding to the diatom *C. neogracile.*

*C. neogracile* is a psychrophilic marine diatom found in Antarctic waters(43). As shown in Figure 5A-D, *C. neogracile* are roughly 10 μm in length with a width of 3-4 μm. Each diatom cell has 1-4 projections protruding from the corners. Given its photosynthetic capability, *C. neogracile* contains chlorophyll that is intrinsically fluorescent. However, the binding of FITC-labelled *Mp*PA14 to *C. neogracile* resulted in a 40-fold increase of fluorescence over the basal autofluorescence of the diatom (Figure 5A, B, Figure S5). The addition of 0.5 mM L-fucose was extremely effective at blocking accumulation of lectin on the diatom (Figure 5C), as the free sugar outcompeted the cell surface glycans for the binding *Mp*PA14 and displaced 95% of the fluorescent signal (Figure 5E). This competitive effect fell off to ~40% as the L-fucose concentration was reduced to 0.1 mM (Figure 5E). The effective concentration of L-fucose needed to block association is significantly higher than the K_d_ of 147 μM for *Mp*PA14-fucose interaction determined by ITC. We reason that glycans coating the diatom cell membrane present numerous end group binding sites in close proximity that can serve as a “molecular velcro” for *Mp*PA14 binding(30).

In contrast to the inhibitory effect of L-fucose, the non-binder of *Mp*PA14, GalNAc, was unable to prevent the lectin from binding the diatom even at 50 mM (Figure 5D, E, Figure S5), validating the results from the binding and structural studies. Importantly, adding 0.5 mM L-fucose to diatoms pre-coated with FITC-labelled *Mp*PA14 resulted in the dissociation of the lectin from the cells. These results suggest that L-fucose can disrupt pre-existing associations between bacteria and diatoms.

## Materials and Methods

### Dextran-based comparative competition assay

The dextran resin competition assay was performed as previously described for *Mh*PA14(12). Briefly, *Mp*PA14 with GFP fused to its N terminus (GFP-*Mp*PA14) was suspended with an aliquot of Superdex-200 (S200) resin. Following an incubation period with gentle mixing, the S200 resin bound with GFP-*Mp*PA14 was pelleted by centrifugation. The pellet was washed twice with 50 mM Tris-HCl (pH 9), 150 mM NaCl and 5 mM CaCl_2_, and the A_280nm_ of the supernatant from the second wash was used as the baseline reading. Next, after resuspension in the same buffer, aliquots of 1.67 μmoles of saccharide were sequentially added to the solution six or seven times with the A_280nm_ of the supernatant being measured after each addition to quantify the release of lectin. The final addition of saccharide was 5 μM. Data from the dextran-affinity assay were plotted using GraphPad Prism after subtracting the background. Next, the data were fitted to a non-linear regression of one-site-specific binding, which follows the model *Y/Bmax* = *X/(K_d_* + *X*), with Bmax as the maximum specific binding and K_d_ as the equilibrium binding constant.

### Isothermal Titration Calorimetry (ITC)

Isothermal calorimetric titration (ITC) measurements were performed at 30 °C with a MicroCal VP-ITC instrument (Malvern). *Mp*PA14 (400 μM) was mixed with serial 5-μl aliquots of 8-mM sugar solution (L-fucose, GlcNAc, or glucose). Sugars were automatically added by a rotating syringe (400 RPM) at 5-min intervals into the *Mp*PA14 solution for a total of 50 injections. The data were analyzed by Origin software Version 5.0 (MicroCal).

### Co-crystallization, X-ray diffraction and structure solutions of MpPA14 with various sugars

Details for the cloning, expression, purification, and crystallization of *Mp*PA14 were previously reported(12, 21). Co-crystallization of *Mp*PA14 with various sugars was performed using the “microbatch-under-oil” method by mixing equal volumes of ~ 20 mg/mL protein with a precipitant solution composed of 0.2 M calcium chloride, 0.1 M HEPES (pH 7), 20% (v/v) polyethylene glycol 3350 and 0.5-1M of different sugars. In addition to the previously reported *Mp*PA14-glucose structure, the 14 different new sugars that were complexed with *Mp*PA14 were: L-fucose, GlcNAc, galactose, allose, mannose, 3-O-methyl-glucose, 2-deoxy-glucose, α-methyl-glucose, myo-inositol, sucrose, trehalose, L-arabinose, ribose, and 2-deoxy-ribose.

X-ray crystallographic data were collected at either the 08ID-1 beamline of the Canadian Light Source synchrotron facility or at the 23-ID-B beamline of the Advanced Photon Source via remote access. Data were indexed and integrated with X-ray Detector Software (XDS)(44) and CCP4-Aimless(45) or the DIALS/xia2 in the CCP4i2 software suite(46). The structure solutions for all complexes were obtained by molecular replacement using the *Mp*PA14 glucose-bound structure as the search model(21). The structures were refined using CCP4-Refmac5(47) or Phenix(48).

### Glycan arrays

Three different glycan arrays were probed with *Mp*PA14. Two of the arrays focused on fungal, bacterial, algal and plant polysaccharides, and the other on mammalian glycans. The first glycan array was done at the Carbohydrate Microarray Facility (Glycosciences Laboratory, Imperial College). GFP-*Mp*PA14 (50 μg/mL) was exposed to the ‘Fungal, bacterial and plant polysaccharide array set 2’, which contained duplicates of 20 saccharide probes from a variety of organisms. An Alexa Fluor 647-tagged anti-GFP antibody was used for detecting the lectin, and the duplicates were averaged to produce the final relative fluorescence unit (RFU) values. In a negative control experiment, where anti-GFP antibody was directly reacted to the saccharide probes, four glycans: lipomannan and lipoarabinomannan from *Mycobacterium tuberculosis,* lipoarabinomannan from *Mycobacterium smegmatis,* and native *o*-glycoprotein from *M. tuberculosis* showed significant binding to the anti-GFP antibody as they produced RFUs of greater than 1,000. This indicated that these four glycan samples yielded false positive results. Therefore, these four polysaccharides were discarded from the analyses shown in the Results section of our paper.

The second glycan array was performed at the Max Planck Institute for Marine Microbiology (Bremen, Germany). The array contained duplicates of 32 polysaccharides including those from macroalgae, bacteria, fungi, and land plants (see details in Table S5). N-terminally His-tagged *Mp*PA14 was incubated with the array and binding of the lectin to the polysaccharides was detected by an anti-His tag secondary antibody conjugated to alkaline phosphatase (Sigma-Aldrich). Microarray probing and quantification were performed as previously described(41).

Maximal mean (average of the duplicates) signal intensity was set to 100 and the rest of values were normalized accordingly. A cut-off of 5 was applied (49).

The third glycan array screening was done by the Consortium for Functional Glycomics (Harvard Medical School) using Version 5.2 of a printed mammalian glycan array, which contained 609 glycans(50). TRIT-C labelled *Mp*PA14 was incubated with the surface-immobilized glycans and the array was scanned at an excitation wavelength of 532 nm. The resulting RFUs were used as a measure of the bound protein. Each glycan was present in six replicates on the array, and the highest and lowest value from each set was omitted to avoid outlying values. The RFU values from the remaining four replicates were averaged.

### Diatom binding experiments

The Antarctic diatom, *Chaetoceros neogracile,* was cultured as previously described(21,43). FITC-labelled *Mp*PA14 (FITC-*Mp*PA14, 0.2 mg/mL) in the presence or absence of sugars was incubated with diatoms in buffer (50 mM Tris-HCl pH 9, 300 mM NaCl, 5 mM CaCl_2_) with gentle mixing for 2 h. Next, diatoms were pelleted by centrifugation for 3 min at 7,000 rpm, and the resulting supernatant was discarded. This procedure was repeated three times to wash away unbound FITC-*Mp*PA14 before the diatom pellet was finally resuspended in 20 μL buffer, which was then used to make slides for fluorescence microscopy. In a separate experiment to test if fucose could compete off the *Mp*PA14 that was already bound to diatoms, FITC-*Mp*PA14 was incubated with diatom for 1.5 h before fucose was added. The rest of the experiment followed the same procedure as described above.

Images were obtained using an Olympus IX83 inverted fluorescence microscope equipped with an Andor Zyla 4.2 Plus camera. Quantification of the fluorescence intensity was done using Fiji ImageJ. The corrected total cell fluorescence (CTCF) was calculated using the formula: CTCF = Integrated Density – (Area of selected cell x Mean Fluorescence of the background)(51). Quantification of 30 individual diatom cells was done for each treatment. Graphs were made using GraphPad Prism.

### Conclusions and Outlook

In this study, we elucidated the molecular basis for ligand recognition by a lectin module widespread in bacterial adhesins. The atomic details revealed by X-ray crystallography not only helped clarify the plasticity of *Mp*PA14 in binding various monosaccharide ligands, but also revealed how the lectin recognizes complex polysaccharides in a more specific manner. We further show that a low millimolar amount of L-fucose can be used to disrupt binding of the lectin to diatom cells. The atomic details for the lectin-carbohydrate interactions elucidated here may serve as the starting points for the development of adhesin antagonists via ligand-based design. For instance, the fact that GlcNAc gains a 2-fold higher affinity for *Mp*PA14 than glucose simply from the replacement of the C-2 hydroxyl with an acetyl group suggests that appending designed substituents on various positions of avid binders such as the C-2 and C-5 of L-fucose might further enhance their potency (Figure 2C, F).

Given the high similarity between *Mp*PA14 and lectin folds in the adhesins of pathogenic bacteria, this work gives insight into how harmful bacterium-host interactions might be controlled through modulation of the lectin-glycan interactions. This anti-adhesion approach holds promise as an alternative or additive approach to treat infections without the excessive use of antibiotics and may thus help mitigate problems with multidrug-resistant bacteria(17, 52, 53).

## General

We are grateful to Dr. John Allingham for the use of his home source X-ray diffractometer at Queen’s University, and to staff members at the Canadian Light Source in Saskatoon and the Advanced Photon Source in Chicago for access to data collection at these synchrotrons. We would like to thank Dr. EonSeon Jin, Hanyang University, Seoul, for the gift of the diatom, *Chaetoceros neogracile* and Dr. Saeed Rismani Yazdi for assistance with the diatom cultures. We thank Dr. Xu Deng from Central South University (China) for fruitful discussions, and for his thoughtful comments as well as those from Drs. Chantelle Capicciotti and Inka Brockhausen on the initial draft of the manuscript. We thank Mr. Kim Munro for acquiring and interpreting ITC data. We appreciate Ms. Sherry Gauthier’s assistance with molecular cloning. In addition, we acknowledge the glycan array facilities at the Consortium for Functional Glycomics (CFG), Harvard Medical School and the Glycosciences Laboratory at Imperial College for their help.

## Funding

This project was funded by a Natural Science and Engineering Research Council (NSERC, http://www.nserc-crsng.gc.ca/index_eng.asp) Discovery Grant (RGPIN-2016-04810). PLD holds the Canadian Research Chair in Protein Engineering, and TDRV held a Canadian Graduate Scholarship from NSERC.

## sAuthor Contributions

S.G. and P.L.D. conceived the study, designed the experiments, and wrote the manuscript. S.G. performed co-crystallization, data collection, and structure determination of the X-ray crystal structures. R.E. and T.D.R.V. performed the comparative binding assays. H.Z. and C.S. performed the diatom binding experiments and analyzed data. S.V. and J.H. performed one of the microbial glycan microarrays and analyzed data. All authors contributed to the revision of drafts of the manuscript.

## Competing interests

The authors declare that they have no competing interests.

## Data and materials availability

X-ray crystal structure coordinates solved in this study have been deposited in the Protein Data Bank with accession codes of 6X7J (*Mp*PA14-fucose), 6X7X (*Mp*PA14-mannose), 6XAQ (*Mp*PA14-α-methyl-glucose), 6X7Z (*Mp*PA14-inositol), 6X7Y (*Mp*PA14-GlcNAc), 6X7T (*Mp*PA14-allose), 6X9M (*Mp*PA14-3-O-methyl-glucose), 6X95 (*Mp*PA14-2-deoxy-glucose), 6XAC (*Mp*PA14-galactose), 6X8D (*Mp*PA14-arabinose), 6X8Y (*Mp*PA14-ribose), 6X9P (*Mp*PA14-2-deoxy-ribose), 6X8A (*Mp*PA14-sucrose) and 6XA5 (*Mp*PA14-trehalose). The data that support the findings of this study are available from the corresponding author P.L.D upon reasonable request.

## Notes

### Competing Interest Statement

The authors have declared no competing interest.

## References

1. A. Varki, Evolutionary Forces Shaping the Golgi Glycosylation Machinery: Why Cell Surface Glycans Are Universal to Living Cells. Csh Perspect Biol 3 (2011).

2. A. Varki, Biological roles of glycans. Glycobiology 27, 3–49 (2017).

3. G. F. Clark, The role of carbohydrate recognition during human sperm-egg binding. Hum Reprod 28, 566–577 (2013).

4. B. Lepenies, R. Lang, Editorial: Lectins and Their Ligands in Shaping Immune Responses. Front Immunol 10 (2019).

5. E. Hebert, Endogenous lectins as cell surface transducers. Bioscience Rep 20, 213–237 (2000).

6. F. S. Ielasi et al., Lectin-Glycan Interaction Network-Based Identification of Host Receptors of Microbial Pathogenic Adhesins. Mbio 7 (2016).

7. D. A. Wesener et al., Recognition of microbial glycans by human intelectin-1. Nat Struct Mol Biol 22, 603–610 (2015).

8. A. B. Boraston, D. N. Bolam, H. J. Gilbert, G. J. Davies, Carbohydrate-binding modules: finetuning polysaccharide recognition. Biochem J 382, 769–781 (2004).

9. Y. van Kooyk, G. A. Rabinovich, Protein-glycan interactions in the control of innate and adaptive immune responses. Nat Immunol 9, 593–601 (2008).

10. M. Maestre-Reyna et al., Structural basis for promiscuity and specificity during Candida glabrata invasion of host epithelia. P Natl Acad Sci USA 109, 16864–16869 (2012).

11. S. Guo, T. D. R. Vance, C. A. Stevens, I. K. Voets, P. L. Davies, RTX Adhesins are Key Bacterial Surface Megaproteins in the Formation of Biofilms. Trends Microbiol 27, 470 (2019).

12. T. D. R. Vance, S. Q. Guo, S. Assaie-Ardakany, B. Conroy, P. L. Davies, Structure and functional analysis of a bacterial adhesin sugar-binding domain (vol 14, e0220045, 2019). Plos One 14 (2019).

13. K. A. Krogfelt, H. Bergmans, P. Klemm, Direct evidence that the FimH protein is the mannose-specific adhesin of Escherichia coli type 1 fimbriae. Infect Immun 58, 1995–1998 (1990).

14. A. Wellens et al., Intervening with urinary tract infections using anti-adhesives based on the crystal structure of the FimH-oligomannose-3 complex. PLoS One 3, e2040 (2008).

15. L. K. Mydock-McGrane, T. J. Hannan, J. W. Janetka, Rational design strategies for FimH antagonists: new drugs on the horizon for urinary tract infection and Crohn’s disease. Expert Opin Drug Discov 12, 711–731 (2017).

16. M. M. Sauer et al., Binding of the Bacterial Adhesin FimH to Its Natural, Multivalent High-Mannose Type Glycan Targets. J Am Chem Soc 141, 936–944 (2019).

17. C. K. Cusumano et al., Treatment and prevention of urinary tract infection with orally active FimH inhibitors. Sci Transl Med 3, 109ra115 (2011).

18. M. Totsika et al., A FimH inhibitor prevents acute bladder infection and treats chronic cystitis caused by multidrug-resistant uropathogenic Escherichia coli ST131. J Infect Dis 208, 921–928 (2013).

19. C. N. Spaulding et al., Selective depletion of uropathogenic E. coli from the gut by a FimH antagonist. Nature 546, 528–532 (2017).

20. S. Guo, C. P. Garnham, J. C. Whitney, L. A. Graham, P. L. Davies, Re-evaluation of a bacterial antifreeze protein as an adhesin with ice-binding activity. PLoS One 7, e48805 (2012).

21. S. Guo et al., Structure of a 1.5-MDa adhesin that binds its Antarctic bacterium to diatoms and ice. Sci Adv 3, e1701440 (2017).

22. S. Guo et al., Conserved structural features anchor biofilm-associated RTX-adhesins to the outer membrane of bacteria. FEBS J 285, 1812–1826 (2018).

23. D. J. Rigden, L. V. Mello, M. Y. Galperin, The PA14 domain, a conserved all-beta domain in bacterial toxins, enzymes, adhesins and signaling molecules. Trends Biochem Sci 29, 335–339 (2004).

24. M. E. Taylor, K. Drickamer, Convergent and divergent mechanisms of sugar recognition across kingdoms. Curr Opin Struct Biol 28, 14–22 (2014).

25. C. Bergmann et al., Spectrum of mutations in the gene for autosomal recessive polycystic kidney disease (ARPKD/PKHD1). J Am Soc Nephrol 14, 76–89 (2003).

26. E. Yoshida et al., Role of a PA14 domain in determining substrate specificity of a glycoside hydrolase family 3 beta-glucosidase from Kluyveromyces marxianus. Biochem J 431, 39–49 (2010).

27. V. V. Zverlov, I. Y. Volkov, T. V. Velikodvorskaya, W. H. Schwarz, Thermotoga neapolitana bglB gene, upstream of lamA, encodes a highly thermostable beta-glucosidase that is a laminaribiase. Microbiology 143 (Pt 11), 3537–3542 (1997).

28. B. P. Cormack, N. Ghori, S. Falkow, An adhesin of the yeast pathogen Candida glabrata mediating adherence to human epithelial cells. Science 285, 578–582 (1999).

29. M. Veelders et al., Structural basis of flocculin-mediated social behavior in yeast. P Natl Acad Sci USA 107, 22511–22516 (2010).

30. L. L. Kiessling, Chemistry-driven glycoscience. Bioorgan Med Chem 26, 5229–5238 (2018).

31. M. Ambrosi, N. R. Cameron, B. G. Davis, Lectins: tools for the molecular understanding of the glycocode. Org Biomol Chem 3, 1593–1608 (2005).

32. N. Garber, U. Guempel, N. Gilboagarber, R. J. Doyle, Specificity of the Fucose-Binding Lectin of Pseudomonas-Aeruginosa. Fems Microbiol Lett 48, 331–334 (1987).

33. R. Loris, D. Tielker, K. E. Jaeger, L. Wyns, Structural basis of carbohydrate recognition by the lectin LecB from Pseudomonas aeruginosa. J Mol Biol 331, 861–870 (2003).

34. D. J. Wilbur, C. Williams, A. Allerhand, Detection of the furanose anomers of D-mannose in aqueous solution. Application of carbon-13 nuclear magnetic resonance spectroscopy at 68 MHz. Journal of the American Chemical Society 99, 5450–5452 (1977).

35. K. H. Ueberschar, E. O. Blachnitzky, G. Kurz, Reaction mechanism of D-galactose dehydrogenases from Pseudomonas saccharophila and Pseudomonas fluorescens. Formation and rearrangemnt of aldono-1,5-lactones. Eur JBiochem 48, 389–405 (1974).

36. S. J. Angyal, The Composition and Conformation of Sugars in Solution. ANGEWANDTE CHEMIEINTERNATIONAL EDITION 8, 157–226 (1969).

37. C. D. Rillahan, J. C. Paulson, Glycan microarrays for decoding the glycome. Annu Rev Biochem 80, 797–823 (2011).

38. S. J. Lim, W. M. Wan Aida, S. Schiehser, T. Rosenau, S. Bohmdorfer, Structural elucidation of fucoidan from Cladosiphon okamuranus (Okinawa mozuku). Food Chem 272, 222–226 (2019).

39. M. T. Ale, A. S. Meyer, Fucoidans from brown seaweeds: an update on structures, extraction techniques and use of enzymes as tools for structural elucidation. RSC Advances 3, 8131–8141 (2013).

40. Z. Li, T. Feizi, The neoglycolipid (NGL) technology-based microarrays and future prospects. FEBS Lett 592, 3976–3991 (2018).

41. S. Vidal-Melgosa et al., A new versatile microarray-based method for high throughput screening of carbohydrate-active enzymes. J Biol Chem 290, 9020–9036 (2015).

42. D. C. Smith, G. F. Steward, R. A. Long, F. Azam, Bacterial mediation of carbon fluxes during a diatom bloom in a mesocosm. Deep Sea Research Part II: Topical Studies in Oceanography 42, 75–97 (1995).

43. I. G. Gwak, W. S. Jung, H. J. Kim, S. H. Kang, E. Jin, Antifreeze Protein in Antarctic Marine Diatom, Chaetoceros neogracile. Mar Biotechnol 12, 630–639 (2010).

44. W. Kabsch, Integration, scaling, space-group assignment and post-refinement. Acta Crystallogr D Biol Crystallogr 66, 133–144 (2010).

45. P. Evans, Scaling and assessment of data quality. Acta Crystallogr D 62, 72–82 (2006).

46. J. Beilsten-Edmands et al., Scaling diffraction data in the DIALS software package: algorithms and new approaches for multi-crystal scaling. Acta Crystallogr D Struct Biol 76, 385–399 (2020).

47. A. A. Vagin et al., REFMAC5 dictionary: organization of prior chemical knowledge and guidelines for its use. Acta Crystallogr D 60, 2184–2195 (2004).

48. P. V. Afonine et al., Towards automated crystallographic structure refinement with phenix.refine. Acta Crystallogr D Biol Crystallogr 68, 352–367 (2012).

49. I. Moller et al., High-throughput mapping of cell-wall polymers within and between plants using novel microarrays. Plant J 50, 1118–1128 (2007).

50. J. Heimburg-Molinaro, X. Song, D. F. Smith, R. D. Cummings, Preparation and analysis of glycan microarrays. Curr Protoc Protein Sci Chapter 12, Unit12 10 (2011).

51. A. Burgess et al., Loss of human Greatwall results in G2 arrest and multiple mitotic defects due to deregulation of the cyclin B-Cdc2/PP2A balance. P Natl Acad Sci USA 107, 12564–12569 (2010).

52. A. M. Krachler, K. Orth, Targeting the bacteria-host interface: Strategies in anti-adhesion therapy. Virulence 4, 284–294 (2013).

53. V. Kalas et al., Structure-based discovery of glycomimetic FmlH ligands as inhibitors of bacterial adhesion during urinary tract infection. Proc Natl Acad Sci U S A 115, E2819–E2828 (2018).

