## supplementary information for "Structural basis of ligand selectivity by a bacterial adhesin lectin involved in multi- species biofilm formation"

### for

### Molecular basis for ligand recognition by a bacterial adhesin lectin involved in multi-species biofilm formation

**Table of contents:**


Figure S1: Structural features of *Mp*PA14.

Figure S2: Structures of *Mp*PA14 in complex with glucopyranose-containing sugars.

Figure S3: Structures of strong-binding, moderate-binding and weak-binding ligands of *Mp*PA14.

Figure S4: Structures of *Mp*PA14 in complex with its moderate- and weak-binding ligands.

Figure S5. Representative images of diatoms with various FITC-*Mp*PA14 and sugar treatments.

Table S1-S3: X-ray crystallographic statistics for *Mp*PA14 in complex with 14 different sugars.

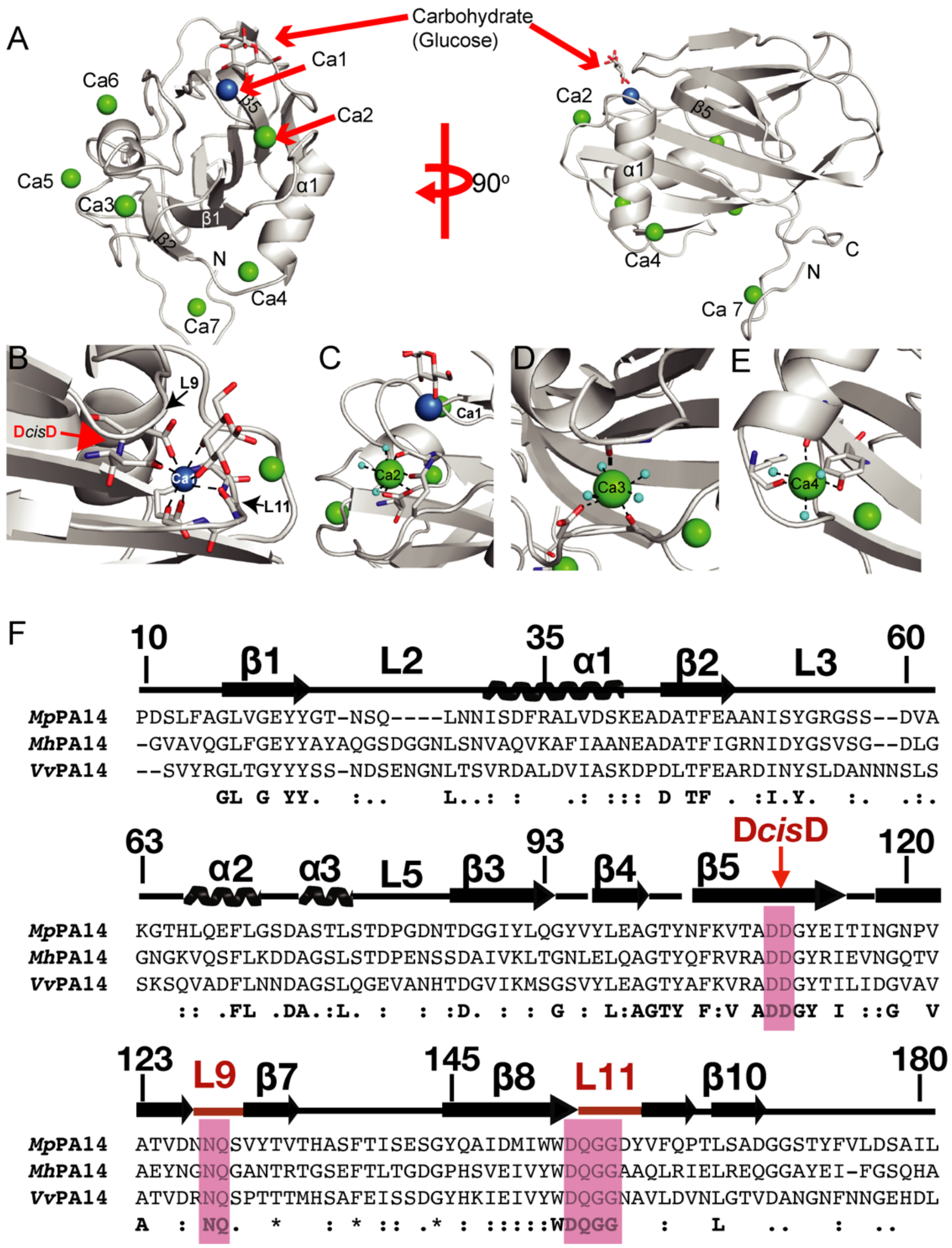


**Figure S1: Structural features of *Mp*PA14.** A) Overview of the *Mp*PA14 structure in complex with glucose. The right panel shows the interface from a view that is rotated approximately 180 ° around a vertical axis from the left. Carbon atoms are colored grey, oxygen atoms are red, and nitrogen atoms are blue. The glucose-coordinationg Ca1 is shown as a dark blue spheres while the other $Ca^{2+}$ ions are shown as green spheres. B) Zoomed-in view of the ligand-binding site of *Mp*PA14. The D*cis*D motif is indicated by an arrow. C)-E) $Ca^{2+}$ coordination sites for Ca2-Ca4. F) Amino-acid alignment of PA14 domains from *M. primoryensis* (*Mp*PA14), the oil-eating bacterium *M. hydrocarbonoclasticus* (*Mh*PA14) and the flesh-eating human pathogen *Vibrio vulnificus* (*Vv*PA14). Amino-acid residue number and secondary structure components for *Mp*PA14 are indicated. Conserved residues that constitute the carbohydrate-binding site are highlighted in magenta. Red arrowhead points to the D*cis*D motif. Loops (L9 and L11) involved in carbohydrate binding are indicated with red color.

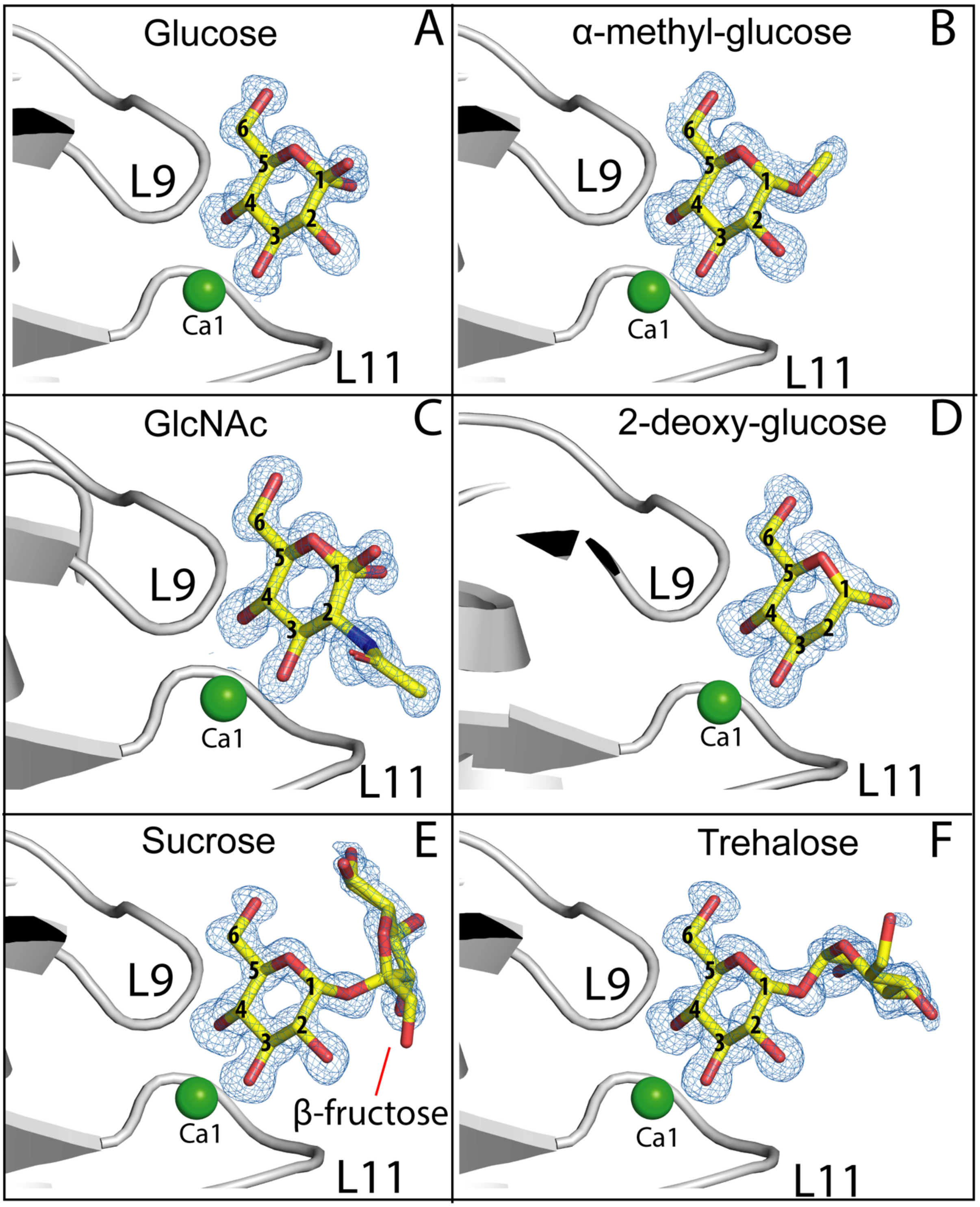


**Figure S2: Structures of *Mp*PA14 in complex with glucopyranose-containing sugars.** The color scheme and labeling are the same as in Fig.2.

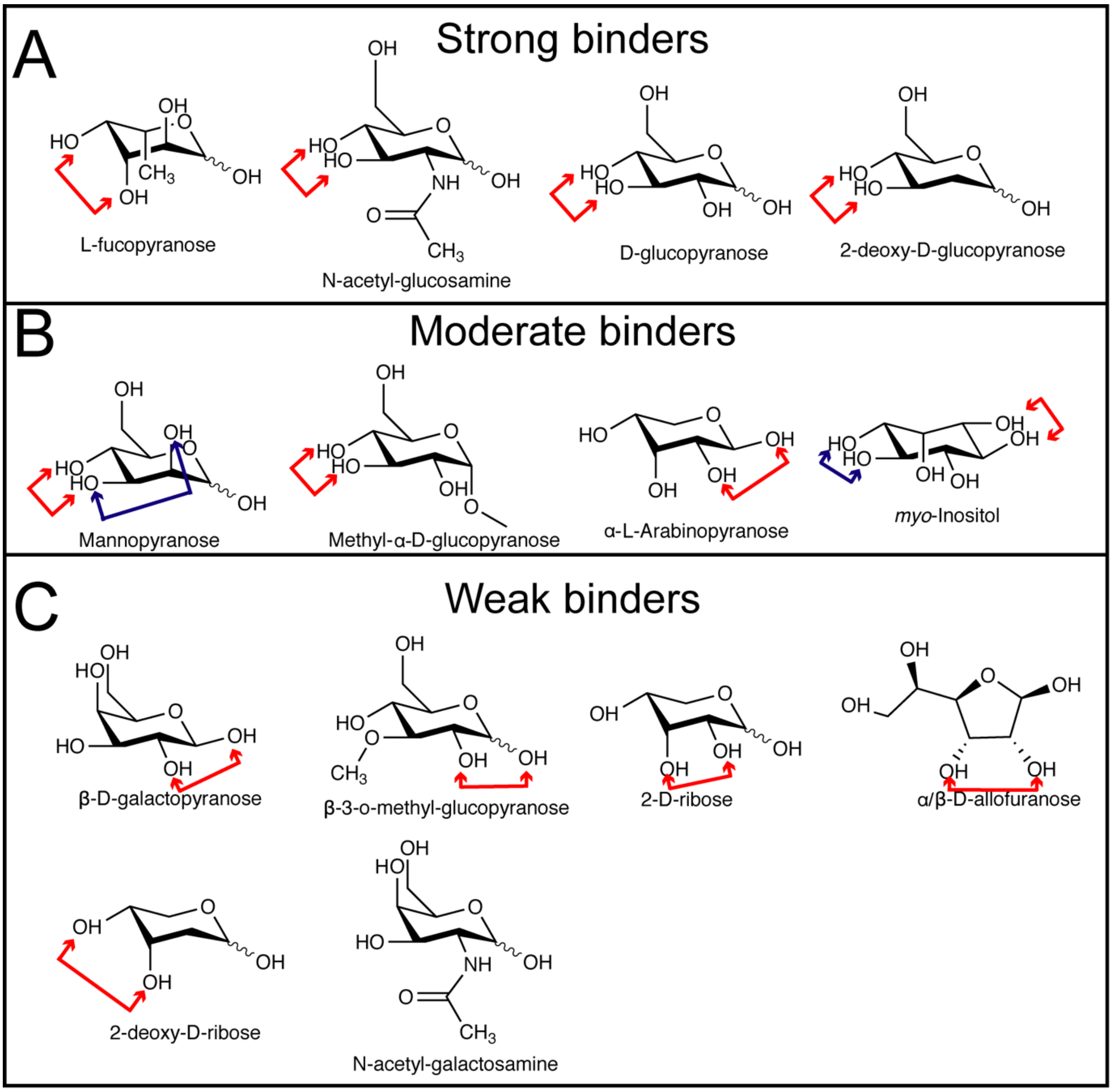


**Figure S3: Structures of strong-binding (A), moderate-binding (B) and weak-binding (C) sugars of *Mp*PA14.** The specific isomer shown for each monosaccharide was revealed by X-ray crystallography as described in Figures 2-3 and S2, 4. Double arrowheads point to the diol groups used by the sugar for interaction with Ca1 in the ligand-binding site of *Mp*PA14. Alternative binding configurations are indicated with double arrowheads in blue or green instead of red.

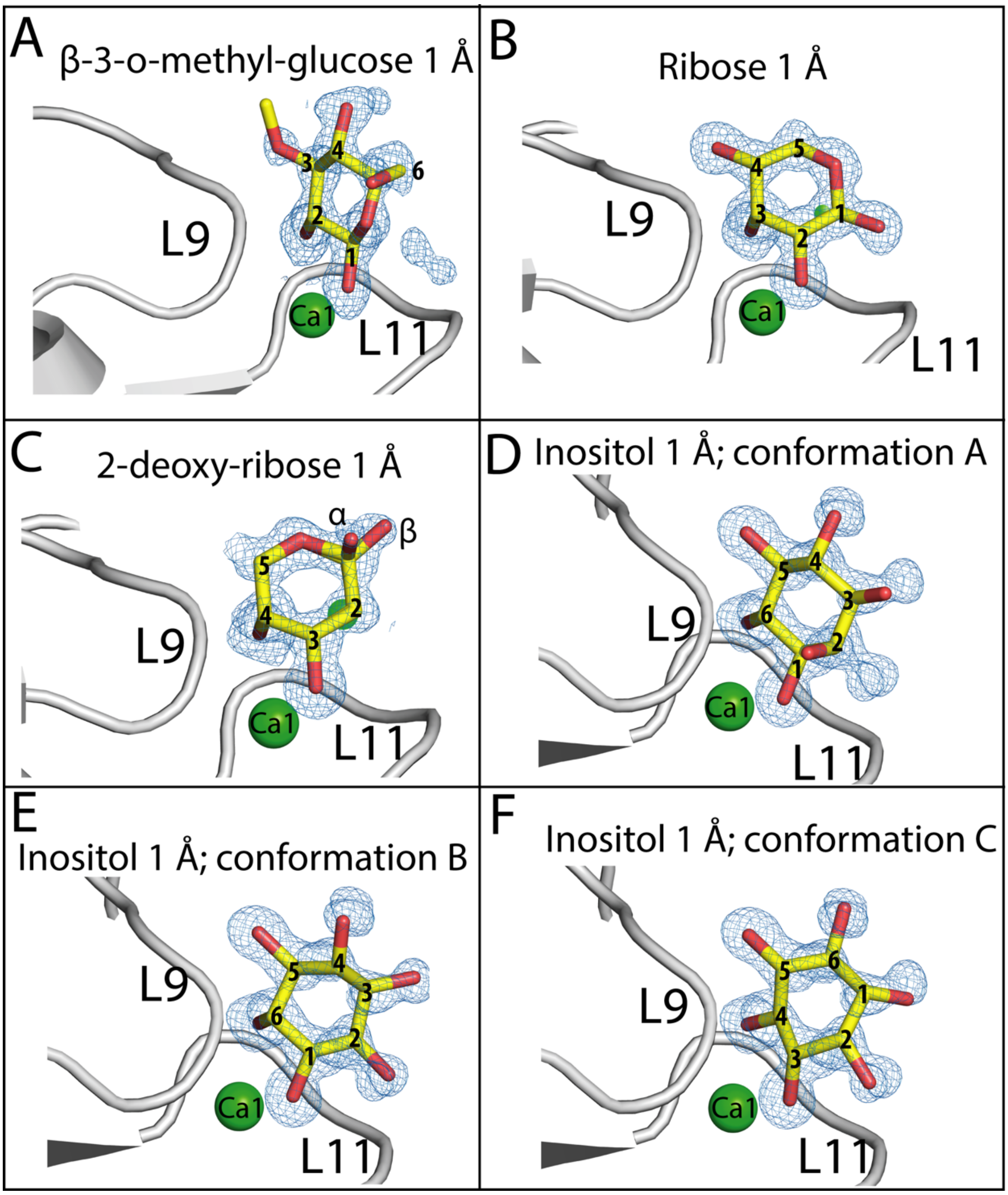


**Figure S4: Structures of *Mp*PA14 in complex with its moderate- and weak-binding ligands.** (A) β-3-o-methyl-glucose. (B) Ribose. (C) 2-deoxy-ribose. (D-F) three binding conformations of Inositol. The color scheme and labeling are the same as in Fig.S2.

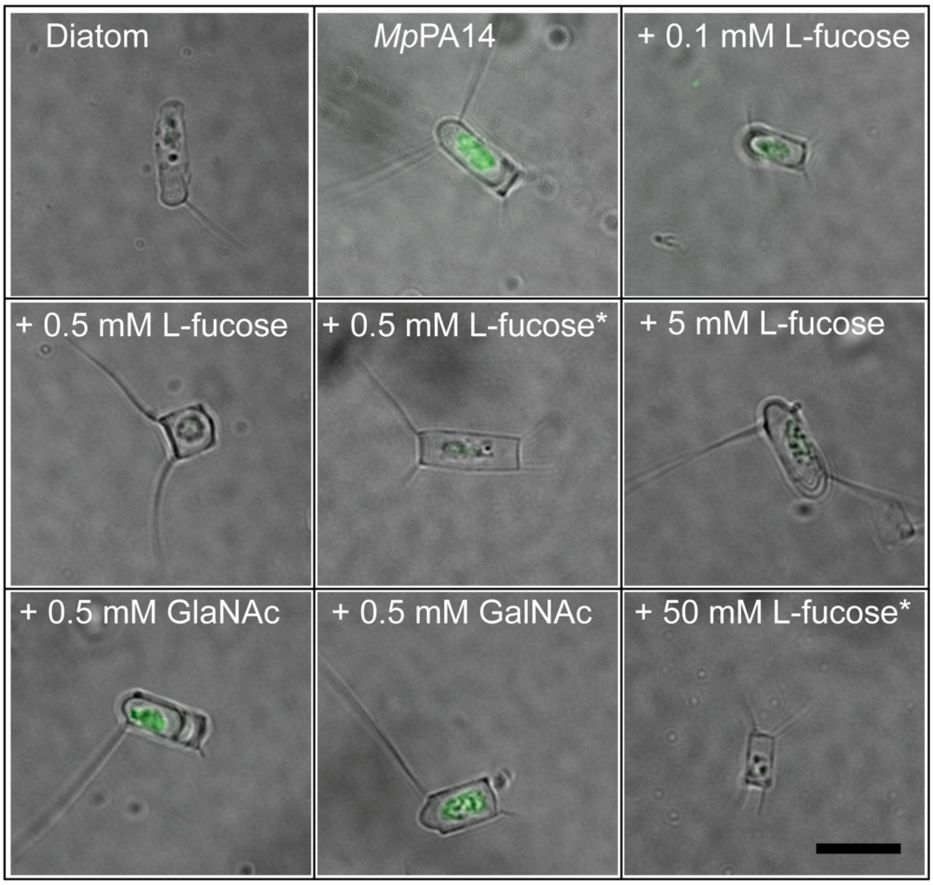


**Figure S5. Representative images of diatoms with various FITC-*Mp*PA14 and sugar treatments shown in Figure 5.** Asterisks (*) represent experiments where diatoms were incubated with FITC-*Mp*PA14 before L-fucose was added.

**Table S1:** X-ray crystallographic statistics for *Mp*PA14 in complex with L-fucose, mannose, α-methyl-Glucose, Inositol and GlcNAc.

| **Data collection** | **L-fucose** | **Mannose** | **α-methyl-Glc** | **Inositol** | **GlcNAc** |
|---|---|---|---|---|---|
| **PDB code** | 6X7J | 6X7J | 6XAQ | 6X7Z | 6X7Y |
| Space group | P 21 21 21 | P 21 21 21 | P 21 21 21 | P 21 21 21 | P 21 21 21 |
| **Cell dimensions** | | | | | |
| (a, b, c) (Å) | 45.08 50.81 79.49 | 45.30, 50.36, 79.13 | 45.12, 50.50, 79.39 | 45.15, 50.57, 79.62 | 45.24 50.65 79.42 |
| (α, β, γ) (°) | 90.0, 90.0, 90.0 | 90.0, 90.0, 90.0 | 90.0, 90.0, 90.0 | 90.0, 90.0, 90.0 | 90.0, 90.0, 90.0 |
| Resolution (Å) | 42.81 - 0.97 | 79.1 - 1.26 | 50.5 - 0.94 | 79.6 - 0.92 | 39.7-0.96 |
| No. of observations | 1175389 | 377043 | 2088235 | 1201596 | 1182800 |
| No. of unique | 100735 | 49044 | 100057 | 111883 | 106951 |
| No. molecules/asymmetric unit | 1 | 1 | 1 | 1 | 1 |
| I/σI | 17.9 (0.8) | 4.79 (1.22) | 17.83 (1.59) | 23.44 (1.63) | 17.1 (1) |
| $R_{merge}$ | 0.076 (1.9) | 0.292 (0.620) | 0.091 (0.901) | 0.057 (0.560) | 0.06(1.03) |
| CC(1/2) | 0.99 (0.36) | 0.95 (0.46) | 0.99 (-0.12) | 0.99 (0.25) | 0.99 (0.49) |
| Completeness (%) | 91.8 (63.7) | 98.8 (83.6) | 84.5 (13.4) | 88.6 (17.2) | 96.1 (70) |
| Multiplicity | 11.7 (5.2) | 7.7 (3.7) | 20.9 (1.7) | 10.7 (1.8) | 11.1 |
| **Refinement** | | | | | |
| Resolution (Å) | 39.7 - 0.97 | 39.6 - 1.3 | 42.6 - 1.2 | 42.7 - 1 | 39.7 - 1 |
| $R_{work}$/ $R_{free}$ (%) | 12.1/13.7 | 15.8/18.6 | 16.3/16.7 | 12.5/13.1 | 12.7/14.1 |
| **No. of atoms** | | | | | |
| protein/Ion/ligand/ water | 2824/7/54/352 | 1576/9/50/265 | 2743/8/27/262 | 2702/6/82/274 | 2666/4/78/252 |
| **B-factors (Å$^2$)** | | | | | |
| protein/Ion/ligand/water | 8.3/8.3/10.1/21.7 | 7.5/13.4/15.2/23.8 | 11.0/12.7/14.8/22.5 | 8.5/9.6/12.6/19.5 | 10.3/9.1/14.2/21.5 |
| **r.m.s deviations** | | | | | |
| Bond lengths (Å) | 0.015 | 0.014 | 0.013 | 0.014 | 0.016 |
| Bond angles (°) | 1.615 | 1.666 | 1.55 | 1.987 | 1.682 |
| **Ramachandron statistics** | | | | | |
| Favored | 96.11 | 94.05 | 95.56 | 96.09 | 96.11 |
| Outliers | 0.56 | 0.54 | 0.56 | 0.56 | 0.56 |

**Table S2:** X-ray crystallographic statistics for *Mp*PA14 in complex with allose, 3-o-methyl glucose, 2-deoxy-glucose, galactose and L-arabinose.

| **Data collection** | Allose | 3-o-methyl glucose | 2-deoxy-glucose | Galactose | L-arabinose |
|---|---|---|---|---|---|
| **PDB code** | 6X7T | 6X9M | 6X95 | 6XAC | 6X8D |
| Space group | P 21 21 21 | P 21 21 21 | P 21 21 21 | P 21 21 21 | P 21 21 21 |
| **Cell dimensions** | | | | | |
| (a, b, c) (Å) | 45.47, 50.27, 79.01 | 45.2, 50.33, 79.14 | 45.19, 50.03, 77.79 | 45.24 50.04 78.71 | 45.12, 50.71, 79.62 |
| (α, β, γ) (°) | 90.0, 90.0, 90.0 | 90.0, 90.0, 90.0 | 90.0, 90.0, 90.0 | 90.0, 90.0, 90.0 | 90.0, 90.0, 90.0 |
| Resolution (Å) | 42.41 - 0.96 | 42.47 - 0.97 | 42.08 - 0.96 | 45.22 - 1.22 | 42.77 - 0.96 |
| No. of observations | 1185822 | 1129177 | 1130090 | 370846 | 1171516 |
| No. of unique | 106122 | 101513 | 103370 | 51321 | 106425 |
| No. molecules/asymmetric unit | 1 | 1 | 1 | 1 | 1 |
| I/σI | 23.1 (2) | 19.9 (1) | 17.1 (0.2) | 28.9 (8.8) | 19.1 (0.9) |
| $R_{merge}$ | 0.058 (0.64) | 0.058 (1.19) | 0.055 (5.8) | 0.044 (0.092) | 0.064 (1.6) |
| CC(1/2) | 0.99 (0.76) | 0.99 (0.57) | 0.99 (0.12) | 0.99 (0.98) | 0.99 (0.62) |
| Completeness (%) | 95.6 (66.2) | 94.6 (59.8) | 96.3 (67.7) | 95.1 (65.9) | 95.2 (62.7) |
| Multiplicity | 11.2 | 11.1 | 10.9 | 7.2 | 11 |
| **Refinement** | | | | | |
| Resolution (Å) | 42.4 - 1 | 42.5 - 1 | 42.1 - 1.05 | 42.2 - 1.22 | 42.8 - 1 |
| $R_{work}$/ $R_{free}$ (%) | 12.7/13.0 | 14.2/15.9 | 16.0/17.3 | 13.4/16.0 | 14.6/16.6 |
| **No. of atoms** | | | | | |
| protein/Ion/ligand/ water | 2772/6/48/282 | 2679/5/73/301 | 2656/5/61/287 | 1439/7/12/255 | 2755/6/50/285 |
| **B-factors (Å$^2$)** | | | | | |
| protein/Ion/ligand/water | 9.6/7.8/21.0/24.7 | 10.9/8.0/20.0/23/5 | 13.0/10.0/30.6/25.1 | 13.7/15.7/14.4/27.0 | 12.8/8.6/24.4/23.2 |
| **r.m.s deviations** | | | | | |
| Bond lengths (Å) | 0.011 | 0.009 | 0.008 | 0.016 | 0.017 |
| Bond angles (°) | 1.45 | 1.321 | 1.173 | 1.689 | 1.672 |
| **Ramachandran statistics** | | | | | |
| Favored | 96.11 | 96.11 | 96.11 | 95.68 | 96.07 |
| Outliers | 0.56 | 0.56 | 0.56 | 1.08 | 0.56 |

**Table S3.** X-ray crystallographic statistics for *Mp*PA14 in complex with ribose, 2-deoxy-ribose, sucrose and trehalose.

| Data collection | Ribose | 2-deoxy ribose | Sucrose | Trehalose |
|---|---|---|---|---|
| **PDB code** | 6X8Y | 6X9P | 6X8A | 6XA5 |
| Space group | P 21 21 21 | P 21 21 21 | P 21 21 21 | P 21 21 21 |
| **Cell dimensions** | | | | |
| (a, b, c) (Å) | 45.14, 50.72, 79.61 | 45.22, 50.53, 79.47 | 45.23 50.61 79.60 | 45.27 50.64 79.43 |
| (α, β, γ) (°) | 90.0, 90.0, 90.0 | 90.0, 90.0, 90.0 | 90.0, 90.0, 90.0 | 90.0, 90.0, 90.0 |
| Resolution (Å) | 42.78 - 0.96 | 39.74 - 0.96 | 42.71 - 1.06 | 42.7 - 1.03 |
| No. of observations | 1152123 | 1182113 | 499930 | 564807 |
| No. of unique | 103936 | 106580 | 61637 | 75689 |
| No. molecules/asymmetric unit | 1 | 1 | 1 | 1 |
| I/σI | 10.1 (0.3) | 17.8 (0.4) | 15.3 (1.2) | 28.0 (2.6) |
| $R_{merge}$ | 0.14 (6.3) | 0.06 (2.8) | 0.11 (1.2) | 0.047 (0.34) |
| CC(1/2) | 0.99 (0.33) | 0.99 (0.12) | 0.99 (0.28) | 0.99 (0.88) |
| Completeness (%) | 92.8 (47.3) | 95.8 (65.0) | 74.6 (11) | 84.2 (15.6) |
| Multiplicity | 1.1 | 11.1 | 8.1 | 7.5 |
| **Refinement** | | | | |
| Resolution (Å) | 42.8 - 1 | 39.7 - 1 | 42.7 - 1.06 | 42.7-1.03 |
| $R_{work}$/ $R_{free}$ (%) | 15.5/16.7 | 12.9/14.2 | 14.2/16.3 | 11.8/13.1 |
| **No. of atoms** | | | | |
| protein/Ion/ligand/ water | 1391/4/22/283 | 2719/5/78/306 | 1432/7/23/265 | 2748/7/45/290 |
| **B-factors (Å$^2$)** | | | | |
| protein/Ion/ligand/water | 8.5/9.1/17.0/23.8 | 11.0/10.3/22.8/25.2 | 9.4/12.0/22.5/24.2 | 9.4/8.7/20.9/20.9 |
| **r.m.s deviations** | | | | |
| Bond lengths (Å) | 0.013 | 0.008 | 0.011 | 0.008 |
| Bond angles (°) | 1.535 | 1.222 | 1.442 | 1.224 |
| **Ramachandran statistics** | | | | |
| Favored | 96.11 | 96.11 | 95.11 | 95.68 |
| Outliers | 0.56 | 0.56 | 1.09 | 1.08 |
